# Evaluation of anti insulin receptor antibodies as potential novel therapies for human insulin receptoropathy

**DOI:** 10.1101/223883

**Authors:** Gemma. V. Brierley, Kenneth. Siddle, Robert. K. Semple

## Abstract

Biallelic loss-of-function mutations in the insulin receptor (INSR) commonly cause extreme insulin resistance and early mortality. Therapeutic options are limited, but anti-INSR antibodies have previously been shown to activate two mutant receptors. This study evaluated four murine anti-INSR monoclonal antibodies, each capable of stimulating wildtype INSR signaling, against ten known pathogenic INSR mutants and one novel mutation, F248C. All the mutant INSR bound antibody. Eight mutants showed antibody-induced autophosphorylation while co-treatment with antibody and insulin increased maximal phosphorylation. After simultaneous knockdown of mouse Insr and expression of mutant INSR in 3T3-L1 adipocytes two antibodies activated signaling *via* AKT preferentially over signaling via ERK1/2 for seven mutants. Two antibodies (83-7 and 83-14) stimulated glucose uptake *via* P193L, S323L, F382V, and D707A mutant INSR, with antibody response greater than insulin response for D707A. These findings suggest that selected monoclonal anti-INSR antibodies elicit sufficient signaling by some mutated INSR to be clinically significant.

**One sentence summary:** Anti-insulin receptor monoclonal antibodies can activate selected naturally occurring mutated human insulin receptors, raising the prospect of novel therapy for lethal recessive insulin resistance.

## Introduction

Insulin downregulates catabolic and activates anabolic pathways, suppresses apoptosis and promotes mitosis in response to nutrient ingestion by activating a homodimeric receptor tyrosine kinase*(1, 2)*. Human loss-of-function mutations in the insulin receptor (INSR) were first reported in 1988*(3, 4)*, since when more than 100 alleles causing severe insulin resistance have been described*(5)*. Biallelic INSR mutations produce extreme congenital insulin resistance, clinically described as Donohue or Rabson Mendenhall syndromes (OMIM#246200 or 262190), both featuring impaired linear growth, soft tissue overgrowth and severe metabolic derangement, with demise usually in the first 3 years of life in Donohue syndrome.

Some *INSR* mutations impair receptor processing and cell surface expression. Many mutations, however, are well expressed, but exhibit impaired insulin binding, impaired signal transduction, perturbed recycling kinetics, or a combination of these*(6)*. It has been suggested that the signaling defect of mutant receptors at the cell surface might be circumvented by binding of anti-receptor antibodies, and proof of this principle was provided for two mutations, one in a cell culture model and one as solubilised receptor*(7, 8)*.

Therapeutic use of antibodies has now become well established. In particular, antibodies have been developed as cancer therapeutics to block receptor signaling *(9)*, although therapeutic antibodies are also increasingly being developed for non-cancer indications*(10)*. Interest in the potential of therapies targeting the INSR has recently rekindled, with inhibitory antibodies in Phase 1 human trials*(11)*, and stimulatory antibodies shown to ameliorate diabetes in rodents*(12–15)* and primates*(16)*.

Given these developments, and the high clinical need in recessive insulin receptoropathy, we assessed the effect of monoclonal anti-INSR antibodies *(17–21)* on signaling by a series of disease-causing mutant INSR in cell-based models. We demonstrate that INSR antibodies bind many mutant receptors, triggering autophosphorylation and downstream signaling. For mutant receptors with selective defects in insulin binding, the signaling response to antibody may exceed that elicited by insulin, suggesting that antibodies might be used therapeutically as surrogate ligands to ameliorate recessive conditions for which there is currently no effective therapy.

## Materials and Methods

### Cell lines and culture conditions

Culture media for CHO Flp-IN cells (Invitrogen) and 3T3-L1 pre-adipocytes (Zenbio) are shown in Table S1. 3T3-L1 pre-adipocytes were grown to confluence and differentiation was induced by Differentiation medium 1 for 72h before changing to Differentiation medium 2 for a further 72h. Adipocytes were maintained in Adipocyte medium containing 1μM insulin ± 1 μg/ml doxycycline. Experiments were undertaken at day 14 or 16 of differentiation as indicated.

### hINSR mutant expression constructs and generation of CHO FlpIN hINSR cells

Mutation numbering refers to mature hINSR ex11+ (GenBank M1005.1), which was amplified from pDNR-Dual using primers incorporating a C-terminus myc tag. Sub-cloning is detailed in Table S2. Mutations were generated with the Quickchange II XL kit (Stratagene). CHO Flp-IN cells were transfected with pCDNA5/FRT/TO/hINSR and pOG44 using Lipofectamine 2000 (Invitrogen). The polyclonal population surviving Hygromycin B selection was used for experiments.

### Lentivirus production and infection of 3T3-L1 preadipocytes

Target sequences, primers, vectors and sub-cloning are detailed in Table S3. Virus containing concatenated miR-shRNAs was packaged and concentrated as described by Shin et al *(22)*. Single cell 3T3-L1 preadipocyte clones were generated by infection with the lowest MOI of virus needed to confer hygromycin B resistance. Several clones per line were characterised for both endogenous *Insr* knockdown and adipocyte differentiation by Oil-Red-O staining*(23)*. For hINSR re-expression studies, 3T3-L1 MmINSRKD cells were infected with virus containing myc-tagged hINSR transgenes, the lowest MOI needed to confer G418 resistance being used to generate polyclonal populations. hINSR expression was confirmed by Sanger sequencing of cDNA.

### Flow cytometry

CHO Flp-IN hINSR cells were blocked by 5%FCS/ FACS buffer (Table S4) before incubation with primary antibodies for 1h at 4°C. Bound antibodies were detected using FITC-conjugated antimouse IgG and a BD FACSCalibur Flow Cytometer (530nm/30nm bandwidth filter, Benton Dickinson). Stacked overlay histograms were visualised with FCS Express 6 Plus (DeNovo Software, USA).

### Receptor Autophosphorylation Assays

CHO Flp-IN hINSR cells were washed twice and serum starved (16h) before stimulation with insulin, antibody or both, for 10min at 37°C/5%CO_2_ and lysis on ice in lysis buffer (Table S4). Receptors were captured overnight at 4°C on anti-myc antibody 9E10-coated white Greiner Lumitrac 600 96-well plates. Phosphotyrosines on immunocaptured receptors were detected with biotin-conjugated 4G10 platinum phospho-tyrosine antibody and Europium-labelled streptavidin. DELFIA-enhancement solution was added and time-resolved fluorescence measured (Ex:340nm/Em:615nm).

### Downstream signal activation

3T3-L1 adipocytes were washed twice in DMEM, serum starved for 16h in DMEM/0.5%BSA/1μg/ml DOX, and then treated for 10min at 37°C/5%CO_2_ with 10nM insulin, 10nM antibody, or both in DMEM/0.5%BSA. Cells were washed, snap frozen and lysed on ice before two rounds of centrifugation at 16.1x RCF at 4°C for 15min to pellet insoluble material and separate lipids prior to Western blotting.

### Western blotting

10μg of total lysate was resolved on NuPAGE 4–12% bis-tris gels or E-PAGE 48 8% gels (Life Technologies) and transferred to nitrocellulose by iBlot (Life Technologies). Membranes were blocked in 3% BSA/TBST before overnight incubation at 4°C with primary antibodies (Table S5). Horseradish peroxidase (HRP)-conjugated secondary antibodies and Immobilon Western Chemiluminescent HRP substrate (Millipore) were used to detect protein-antibody complexes, and grey-scale 16-bit TIFs captured with an ImageQuant LAS4000 camera system (GE). Each immunoblot in Figures 4 and S2 contained a sample of 3T3-L1 MmINSRKD hINSR WT treated with 10nM insulin.

### Western Blot Image Densitometry

Pixel density of grey-scale 16-bit TIFs was determined in ImageJ 1.47v (NIH, USA). The rectangle tool was used to select lanes and the line tool to enclose the peak of interest and subtract background. The magic-wand tool was used to select the peak area and obtain the raw densitometry value. Mean band intensity of total INSRβ, myc-tagged INSRβ, AKT, ERK1/2, GSK3α/β, p70S6K, and calnexin was used to normalise raw densitometry values for phospho-INSRβ, pAKT, pERK1/2, pGSK3α, pp70S6K, and pAS160. Normalised values for phosphorylated targets were scaled to the mean WT INSR response to insulin.

### Glucose Uptake

3T3-L1 adipocytes were washed twice (DMEM), serum-starved for 16h in low-glucose DMEM/0.2% BSA/1μg/ml DOX, washed twice in PBS and then stimulated for 30min at 37°C/5%CO_2_ with 10nM insulin, 10nM antibody or both in KRPH/0.2%BSA buffer (Table S4). Cells were incubated with 1mM 2-Deoxy-d-glucose for 5min at 37°C/5%CO_2_ before washing (PBS), lysing with 0.1M NaOH, and snap freezing. Glucose uptake was measured by the fluorescence method of Yamamoto et al *(24)*.

### Statistical Analysis

One-way ANOVAs with Tukey’s multiple comparisons test were performed with GraphPad Prism 6 (GraphPad Software). Error bars represent SEM or SD as indicated. All experiments were performed at least three times.

## Results

### Assessment of mutant INSR cell surface expression and antibody binding

Eleven INSR mutations were selected for study (Table S6). Eight were based on evidence of cell surface expression, prioritizing mutations identified in multiple reports to maximize likely availability of patients for future trials. A previously unpublished F248C mutation, identified in a child with Rabson Mendenhall Syndrome, was included opportunistically. The well-studied P1178L tyrosine kinase mutation *(25, 26)*, and the L62P mutation which severely impairs processing*(27)*, were added as controls. Figure 1A displays the extracellular INSR mutations mapped onto the crystal structure of the INSR*(28)*. Four mouse monoclonal anti-human INSR antibodies were used, which had previously all been shown to have partial agonist activity at wild type (WT) receptors, but different effects on kinetics and affinity of insulin binding (Table 1). As Fab fragments of 83-7 and 83-14 were used in determining the crystal structure of insulin-bound INSR*(29)* and their binding epitopes are known and are displayed in Figure 1B.

**Fig 1.**
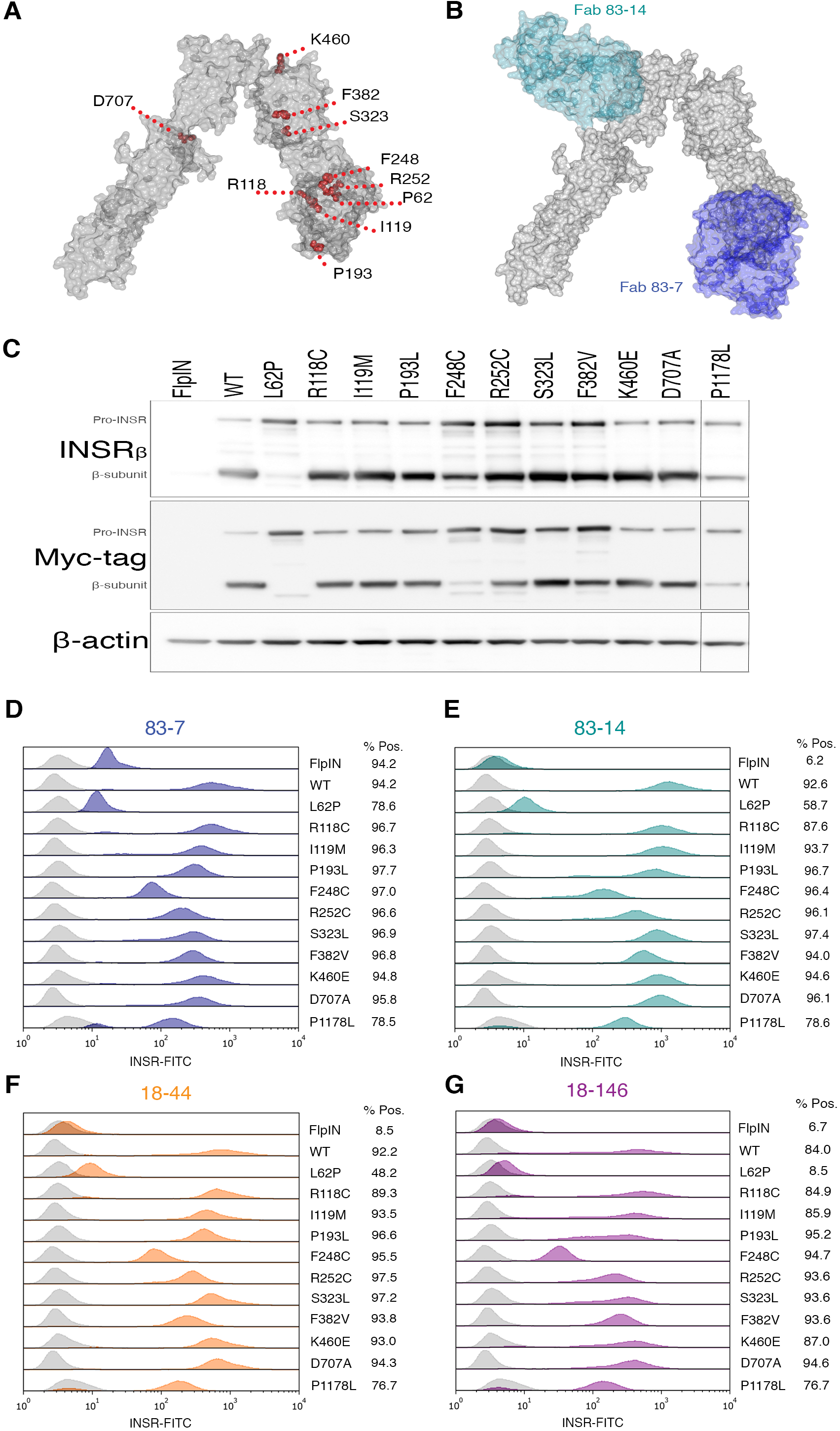
Mutant INSR are expressed at the cell surface and are bound by anti-INSR antibodies. (**A**) INSR monomer PDB structure *4ZXB(22)* visualised with CCP4MG (v. 2.10.6), locations of residues that are mutated in this study are highlighted in red. (**B**) INSR monomer in complex with Fab fragments 83–7 and 83–14, PDB structure 4ZXB. (**C**) Western blot of lysates from CHO Flp-In cells stably expressing human WT or mutant INSR as indicated. In INSR β-subunit and myc-tag blots upper bands are pro-INSR and lower bands are mature processed β-subunits, as indicated. (**D-G**) Cell surface expression of INSR mutants as determined by flow cytometry. Stacked overlay single parameter histograms displaying intensity of INSR-FITC fluorescence on the X-axis and number of events on the Y-axis. Isotype control IgG (light grey) was used as a negative control to generate a negative gate to determine the percentage of the population positive for anti-INSR antibody binding (numbers). Rightward shift of the peak (blue 83–7, dark cyan 83–14, orange 18–44, purple 18–146) from the IgG control is a function of both mutant INSR expression and antibody affinity.

**Table 1.** Characteristics of INSR antibodies studied

|  | **83-7** | **83-14** | **18-44** | **18-146** |
| --- | --- | --- | --- | --- |
| **Type** | *Mm* mAb IgG1 (bivalent) (18) | *Mm* mAb IgG2a (bivalent)(18) | *Mm* mAb IgG2b (bivalent) (18) | *Mm* mAb IgG1 (bivalent) (18) |
| **Epitope** | CR domain* (29) | Fn<sup>I</sup> domain* (29) | Within extracellular β-subunit (18) | Within α-subunit |
| **Effect on INSR:** |  |  |  |  |
| Insulin binding | Modestly enhances (18, 20) | Inhibits (18) | Modestly inhibits (18) | Enhances (18) |
| Pre-bound insulin | No effect on dissociation (19) | Inhibits dissociation (19) | No effect on dissociation (19) | Inhibits dissociation (19) |
| Autophosphorylation | Stimulates (20) | Stimulates (17) | Stimulates (20) | Not assessed |
| Kinase activity | Stimulates (19, 20) | Stimulates (19, 20) | Stimulates (19, 20) | Stimulates (19) |
| **Biological outcomes:** |  |  |  |  |
| Glucose-uptake | Stimulates (17) | Stimulates (17) | Stimulates (17) | Not assessed |
| Lipogenesis | Stimulates (21) | Stimulates (21) | Stimulates (21) | Not assessed |

Type, epitope, and effect on WT INSR insulin binding, pre-bound insulin, autophosphorylation, kinase activity, glucose-uptake and lipogenesis of murine (*Mm*) monoclonal antibodies (mAb) used throughout this study. \*, precise epitope known; CR, cys-rich domain; Fn<sup>I</sup>, first Fibronectin type III domain.

Mutations were introduced into the B isoform of the INSR, believed to be the more important isoform for insulin’s metabolic actions*(30)*. To enable discrimination of endogenous INSR and human INSR mutants a C-terminal myc-tag was used in the mutant constructs. Tagged mutants were expressed in Chinese hamster ovary (CHO) cells using the Flp-In system, ensuring differences in protein expression are due to differential processing or stability of receptor protein rather than differential mRNA expression. The mutants were well processed to mature β-subunits with the exception of L62P, for which beta subunit was barely detectable. More modest reductions were seen for the previously unstudied F248C and for the P1178L mutation (Figure 1C).

Cell surface expression and antibody binding of mutant INSR was assessed by flow cytometry (Figure 1D-G). All INSR antibodies bound each mutant INSR, as shown by right-shifted peaks relative to control IgG, indicating no gross changes in receptor morphology. Poor expression of L62P was in keeping with prior reports*(31)* and L62P was not studied further. The rightward shift for mutants corresponded to expression of mature β-subunits seen by immunoblotting, suggesting that relative shifts reflected differences in receptor expression rather than antibody affinities. Although some mutations are close to the epitope for antibody 83–7, none of the affected residues provide critical antibody contacts. Indeed, no difference in binding of 83-7 to the mutant panel was seen compared to 83-14, which binds to a surface unaffected by the mutations (Figure 1A & B). Antibody 83-7 demonstrated cross-reactivity with endogenous CHO INSR as evidenced by positive staining of CHO Flp-IN parent cells, while the other antibodies did not detectably cross-react.

### Assessment of mutant INSR autophosphorylation in response to antibody and/or insulin

Trans-autophosphorylation of tyrosines in the intracellular INSR is the first detectable signaling event after insulin binding, so the ability of insulin and antibodies to induce tyrosine phosphorylation of mutant INSR was next examined using anti-myc immunoprecipitation and Europium-based immunoassay. Most mutant receptors (P193L, F248C, R252C, S323L, F382V, D707A, P1178L) demonstrated diminished maximal autophosphorylation response to insulin, ranging from 0 – 27% of WT (Figure 2, Table S7, data not shown for non-responsive P1178L). R118C, I119M, and K460E however showed autophosphorylation comparable to WT and so were not studied further. Altered insulin EC50 was discernible only for S323L (Table S7), although the insulin concentration range tested and the small magnitude of responses precluded precise determinations.

**Fig 2.**
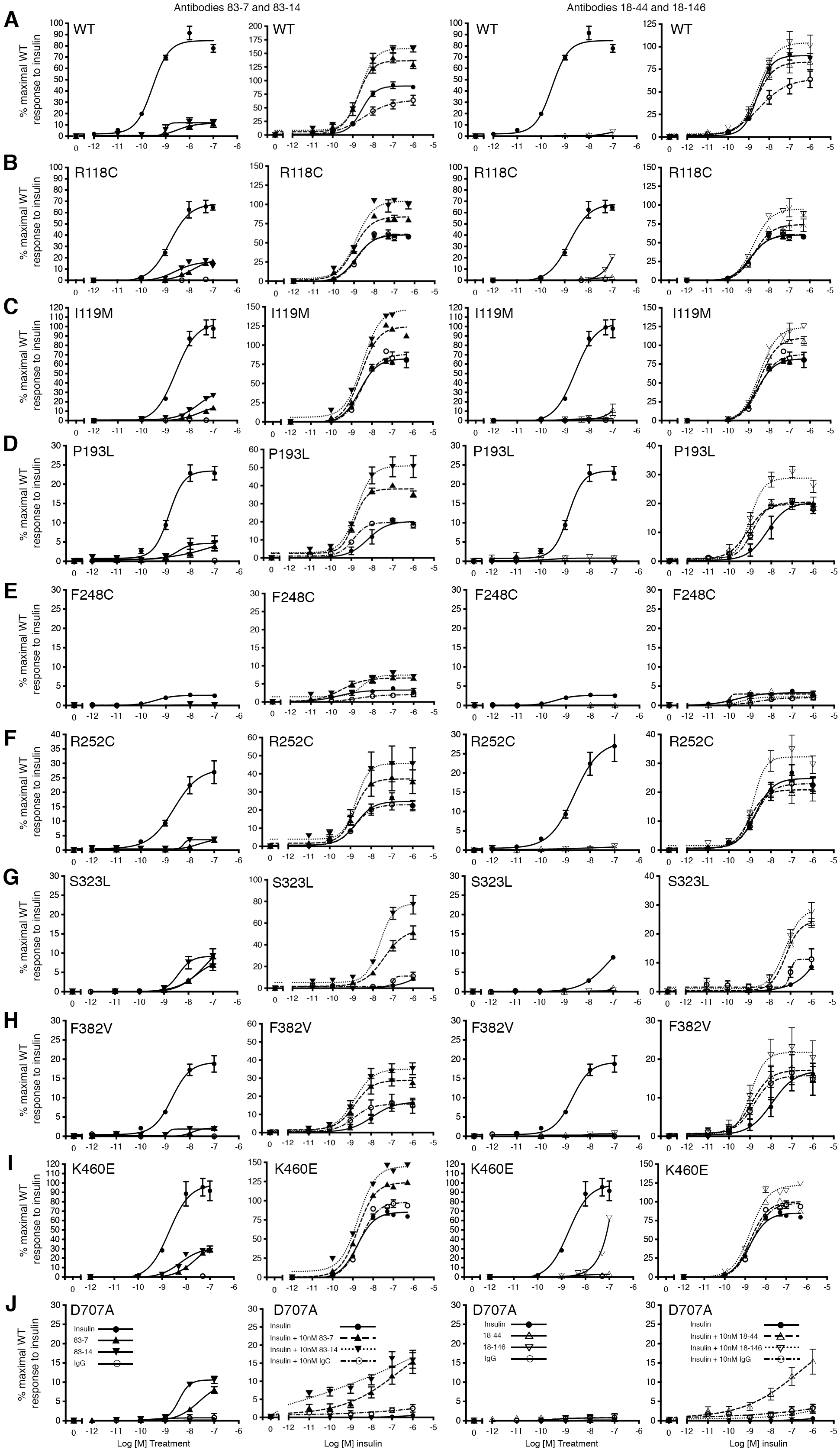
Insulin and antibody stimulated autophosphorylation of WT and mutant INSR. CHO FlpIn cells stably expressing either human WT or mutant INSR (as indicated) were serum starved prior to 10 minute stimulation with increasing concentrations of either insulin, antibody (83-7, 83-14, 18–44, 18–146 or control IgG, solid lines) or increasing concentrations of insulin in the presence of 10nM antibody (broken lines). Cells were lysed and myc-tagged receptors were immunocaptured on 96-well plates and then incubated with biotin-conjugated 4G10 platinum antibody to detect phosphorylated tyrosine residues. Europium-labelled strepavidin was used to detect bound anti-phosphotyrosine antibody 4G10 by time-resolved fluorescence. The data points are the mean ± SEM of duplicate samples from three independent experiments. Error bars are shown when larger than the size of the symbols. In (**A - J**) treatments containing antibodies 83-7 and 83-14 are to the left of the figure, treatments containing antibodies 18–44 and 18–146 are to the right of the figure. Single treatments are denoted by: insulin [solid line •], 83-7 [solid line ▴], 83-14 [solid line ▾], control IgG [solid line ○]. Dual treatments are denoted by: insulin + 10nM 83-7 [dashed line ▴], insulin + 10nM 83-14 [dotted line▾], insulin + 10nM 18–44 [dashed line ▿], insulin + 10nM 18146 [dotted line ▵], insulin + 10nM control IgG [dash-dot line ○]. EC_50_ values are presented in Table S7.

Antibodies 83-7 and 83-14 alone also elicited autophosphorylation of WT and all mutant INSRs except F248C and P1178L; antibodies 18–146 and 18–44 were less effective but also elicited autophosphorylation of WT and several mutant INSRs. In most cases antibody response was lower than insulin response (Figure 2, Table S7), however for S323L the maximal autophosphorylation response to 83-7 and 83-14 was similar to that with insulin, while D707A was activated by antibodies but not insulin.

We next evaluated responses to insulin plus 10nM antibody, based on evidence suggesting this concentration elicits the maximal response*(19, 21)*. In the presence of antibodies 83-7 and 83-14 the maximal response of WT and mutant INSRs to insulin was increased without affecting potency (although EC50 values were not precisely determined) (Figure 2, Table S7). This was observed across all mutant receptors except the kinase-dead P1178L*(25, 26)*. Again, antibodies 18–44 and 18–146 elicited smaller effects.

### Generation of a novel adipocyte cell model of insulin receptoropathy

To assess antibody-induced signaling downstream from the INSR, an adipocyte model of insulin receptoropathy was generated. A Tet-responsive miR-shRNA selectively targeting murine Insr was transduced into 3T3-L1 pre-adipocytes to generate a stable clone (Figure 3A). This was transduced with lentiviruses encoding C-terminal myc-tagged WT or mutant hINSR, also controled by Tet-responsive elements (Figure 3B), generating cells in which doxycycline simultaneously knocked down endogenous murine insulin receptor and induced overexpression of myc-tagged human INSR. This system permitted pre-adipocyte differentiation uncompromised by mutant receptor expression before induction of murine *Insr* knock-down and hINSR re-expression in mature adipocytes (Figure 3C & D). The doxycycline concentration producing maximal *Insr* knockdown resulted in overexpression of hINSR transgenes (Figure 3C and E), however the receptor processing defects observed in CHO cells (Figure 1D) were preserved. The C-terminal myc-tag enabled discrimination of endogenous mouse and ectopic human INSR by size shift of the INSR β subunit on immunoblotting, or by anti-myc antibodies (Figure 3E). The preadipocyte cell lines generated differentiated efficiently into mature adipocytes as evidenced by Oil-red-O staining (Figure 3F).

**Fig 3.**
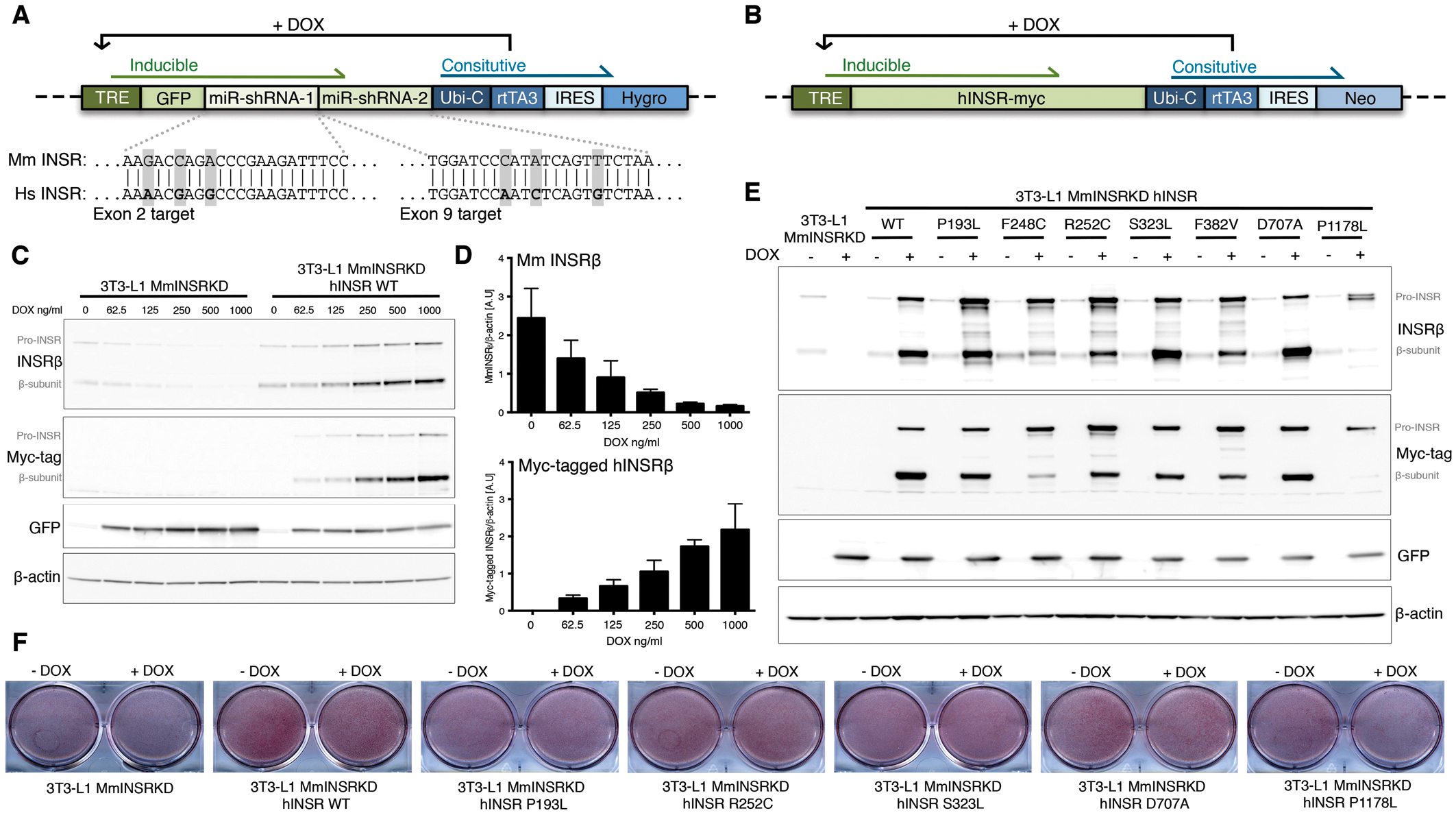
Generation of a novel stable 3T3-L1 adipocyte model of insulin receptoropathy. Concatenated miR-shRNAs targeting murine INSR in exon 2 and exon 9 proceeded by GFP under the control of a Tet-responsive element (**A**) was packaged into third-generation lentivirus to enable transduction of 3T3-L1 preadipocytes. Exploded view shows the nucleotide mismatches between the mouse INSR targeted by each miR-shRNA with the human INSR sequence. Green shaded elements of the transgene are inducible by the addition of doxycycline (DOX). Transduced 3T3-L1 preadipocytes underwent single cell clonal selection in the presence of hygromycin to generate 3T3-L1 MmINSRKD. 3T3-L1 MmINSRKD cells were then transduced with a second lentivirus encoding C-terminal myc-tagged human INSR transgenes under the control of a Tet-responsive element (**B**) and underwent polyclonal selection in the presence of neomycin to generate 3T3-L1 MmINSR KD hINSR. (**C**) Western blots of whole cell lysates from day 10 mature 3T3-L1 MmINSRKD and 3T3-L1 MmINSRKD hINSR WT cells grown in the presence of increasing concentrations of DOX for 72hrs. (**D**) Densitometry analysis of Western blots from three independent experiments demonstrating knockdown of endogenous mouse INSR and expression of human INSR with increasing concentrations of DOX. (**E**) Western blots of whole cell lysates from day 16 mature 3T3-L1 MmINSRKD and 3T3-L1 MmINSRKD hINSR (mutant INSR as indicated) grown in the presence of 1μg/ml DOX for 10 days. (**F**) Oil-red-O staining of lipid accumulation in day 10 mature 3T3-L1 MmINSRKD and 3T3-L1 MmINSRKD hINSR WT or mutant (as indicated) cells grown in ± DOX for 72hrs. Abbreviations: TRE, tet-response element; GFP, green fluorescent protein; Ubi-C, ubiquitin C promoter; rtTA3, reverse tetracycline controlled transactivator; IRES, internal ribosome entry site; Hygro, hygromycin resistance; Neo, neomycin resistance.

### Activation of signaling downstream from mutant INSRs by insulin and antibody

Plasma insulin concentration in human insulin receptoropathies usually lies between 0.3 and 3 nM in the fasting state (Table S6), and at least an order of magnitude higher when fed. We focused on an insulin concentration of 10nM, mimicking the fed disease state. WT INSR autophosphorylation was strongly induced by insulin, but was undetectable after receptor knockdown alone (Figure 4). Otherwise the pattern of autophosphorylation of overexpressed receptors in response to insulin and/or antibody was similar to that seen in CHO cells. Thus, antibodies 83-7 and 83-14 alone induced WT receptor autophosphorylation on Y1162/Y1163, while antibodies 18–44 and 18–146 were less effective. Insulin-stimulated autophosphorylation was reduced by 75–100% in mutant INSRs compared to WT. Although antibodies alone induced low-level phosphorylation of mutant INSRs (<10%), for S323L and D707A responses to antibodies 83-7 and 83-14 were equal to or greater than those with insulin. Combined insulin and antibody treatment enhanced phosphorylation of each of the mutant INSR in the case of 83-7 and 83-14, likely synergistically, but not 18–44 and 18–146 (Figure 4).

**Fig 4.**
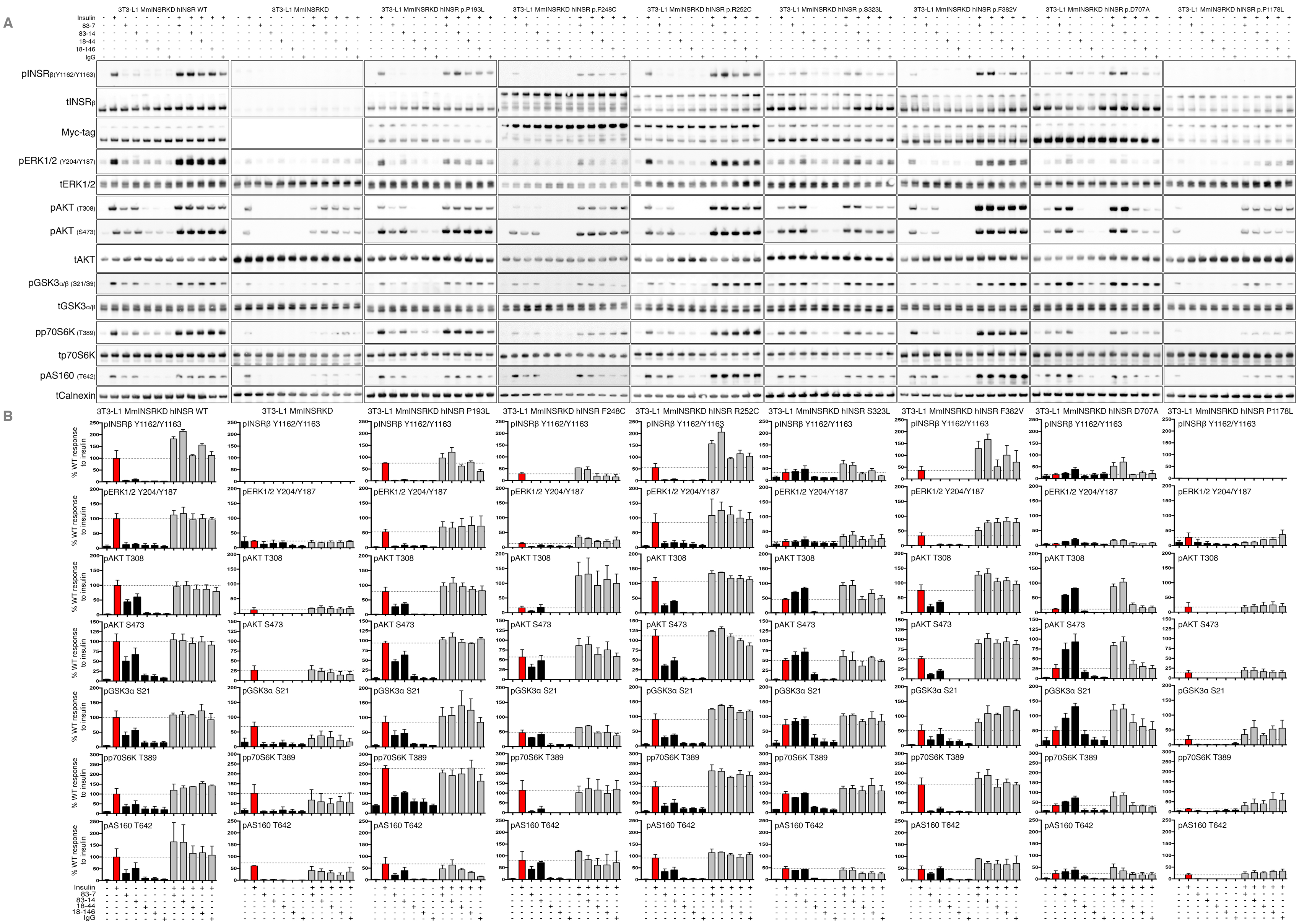
Activation of signaling pathways downstream of WT and mutant INSR by insulin and antibody stimulation. 3T3-L1 MmINSRKD hINSR (mutant as indicated) adipocytes were grown in the presence of 1μg/ml DOX for 8 days prior to overnight serum-starvation on day 13 of differentiation. Adipocytes were then stimulated with either 10nM insulin, 10nM antibody (83-7, 83-14, or control IgG) or 10nM insulin containing 10nM antibody for 10 minutes at 37°C/5%CO_2_. Following stimulation, cells were washed and snap frozen prior to lysis and Western blot (**A**). Phospho-INSRβ, pERK1/2, pAKT, pGSK3α, pp70S6K, and pAS160 densitometry after normalisation to the mean band intensity of total INSRβ, myc-tagged INSRβ, AKT, ERK1/2, GSK3α/β, p70S6K, and calnexin for each biological replicate (**B**). Data are the mean ± SEM of three independent experiments and are expressed as relative to hINSR WT response to insulin stimulation.

AKT2/PKBβ is the critical transducer of the metabolic actions of insulin, being activated by phosphorylation of T308 and S473. Activated AKT2 phosphorylates substrates including glycogen synthase kinase (GSK3α/β), which regulates glycogen synthesis, p70 S6 Kinase (p70S6K), which stimulates protein synthesis, and AS160, which encodes a GTPase-activating protein that restrains GLUT4 vesicle translocation until phosphorylated. Insulin treatment of WT INSR induced strong AKT phosphorylation at both sites, and this was severely attenuated though not abolished by knockdown of endogenous insulin receptor or by knockdown with re-expression of the kinase-dead P1178L mutant (Figure 4). Attenuation of signaling was also apparent downstream of AKT, with phosphorylation of p70S6K and GSK3 only modestly impaired, and AS160 phosphorylation unaffected.

Across the panel of mutants studied several patterns were seen. In D707A receptor-expressing cells, phosphorylation of AKT and its substrates was severely attenuated in response to insulin, while for S323L, F382V, and F248C mutants progressive “escape” from signaling impairment was seen, with lesser impairment of AKT phosphorylation than of receptor autophosphorylation, and only partial inhibition at downstream substrates. P193L and R252C demonstrated similar insulin-induced AKT and AKT substrate phosphorylation to WT receptor.

Antibodies alone stimulated AKT and AKT substrate phosphorylation in all cells except those overexpressing the kinase-dead P1178L mutant. For S323L and D707A mutants antibodies 83-7 and 83-14 stimulated greater phosphorylation than insulin alone, by virtue of the low response of those mutants to insulin, while antibodies 18–44 and 18–146 stimulated only modest phosphorylation (Figure 4). Co-treatment of cells with insulin and antibodies 83-7 and 83-14 enhanced AKT and AKT substrate phosphorylation with respect to insulin alone, without evidence of synergy. Additivity following insulin and antibody co-stimulation was generally less than observed for receptor autophosphorylation in CHO cells (Figure 2).

Activation of the INSR by insulin stimulates not only PI3K/AKT, but also RAS/RAF/MEK/ERK signaling, through both IRS-dependent and IRS-independent mechanisms*(32)*. Activation of this pathway is seen as a surrogate for mitogenicity of insulin analogues*(33)*, which is important in view of concerns about long-term cancer risks of analogues with pro-proliferative activity. Insulin treatment of WT INSR induced robust phosphorylation of ERK1/2 at Y204/Y187, with each mutant INSR displaying reduced phosphorylation in response to insulin compared to WT (Figure 4). Antibody treatment of mutant or WT INSR did not appreciably induce ERK1/2 phosphorylation, while dual stimulation with antibody plus insulin did not increase ERK1/2 phosphorylation compared to insulin alone. Higher basal ERK1/2 phosphorylation was observed in cells with Insr knockdown alone, or with Insr knockdown and P1178L receptor overexpression, but this did not change with any treatment.

### Effect of Insulin and/or antibody on glucose uptake

Stimulation of glucose uptake is a key outcome of INSR activation, and this was assessed in the 3T3-L1 model. Parent 3T3-L1 cells and cells harboring the Insr knockdown construct but not treated with DOX displayed similar high levels of insulin-stimulated glucose-uptake (Figure S1A, B), but insulin did not stimulate uptake in conditional Insr knockdown cells treated with doxycycline (Figure 5). Cells with endogenous mouse Insr knockdown and WT hINSR re-expression, in contrast, demonstrated only a 1.8 fold increase in glucose uptake upon insulin stimulation (Figure S1A). The apparently poor response to insulin was due to increased basal glucose uptake in WT receptor-overexpressing cells (Figure S1B). Basal uptake significantly differed among mutant receptor-expressing cell lines, appearing to reflect mutant receptor function (Figure S1C).

**Fig 5.**
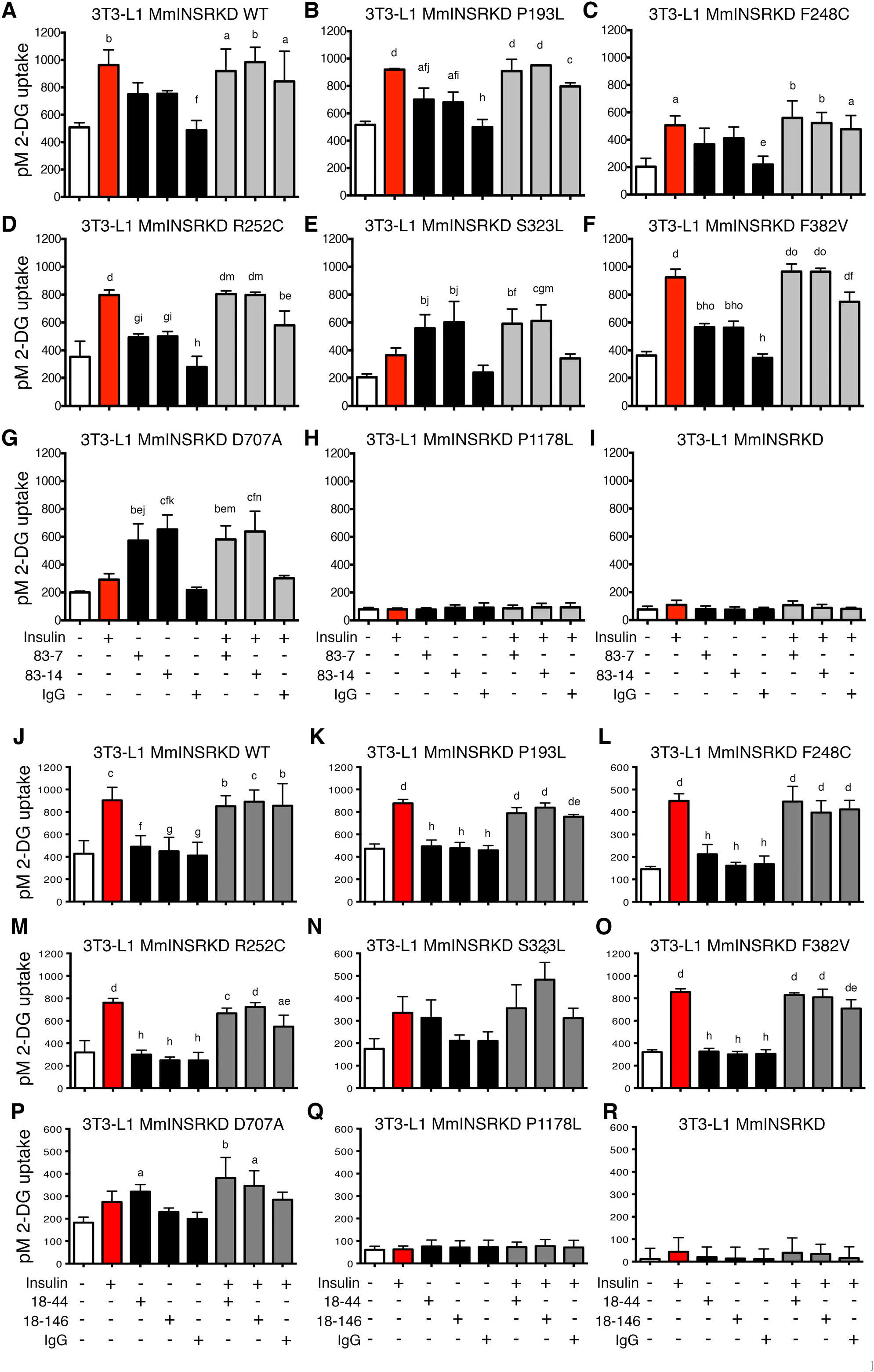
Insulin and antibody stimulated glucose uptake via WT and mutant INSR. 3T3-L1 MmINSRKD hINSR (mutant as indicated) adipocytes were grown in the presence of 1μg/ml DOX for 10 days prior to overnight serum-starvation on day 15 of differentiation. The cells were stimulated for 30 minutes with either 10nM insulin, 10nM antibody (83-7, 83-14, 18–44, 18–146, or control IgG) or 10nM insulin containing 10nM antibody prior to the addition of 2-Deoxy-D-glucose for 5 minutes. Cells were then washed, lysed and assessed for 2-Deoxy-D-glucose uptake. (**A – I**) Glucose uptake stimulated by treatment with antibodies 83-7 and 83-14. (**J – R**) Glucose uptake stimulated by treatment with antibodies 18–44 and 18–146. Data is the mean ± SD from three independent experiments, statistical significance determined by one-way-ANOVA with Tukey’s multiple comparison test. Statistical significance from un-stimulated basal is denoted by a, b, c, d (p<0.05, p<0.01, p<0.001, p<0.0001 respectively). Statistical significance from 10nM insulin treatment is denoted by e, f, g, h (p<0.05, p<0.01, p<0.001, p<0.0001 respectively). Statistical significance from 10nM IgG control treatment is denoted by i, j, k, l. Statistical significance from 10nM insulin in the presence of 10nM IgG control is denoted by m, n, o (p<0.05, p<0.01, p<0.001 respectively).

Despite this loss of dynamic range in the assay, insulin stimulated glucose uptake *via* WT, P193L, F248C, R252C, and F382V receptors (Figure 5). Insulin-stimulated uptake was similar in cells expressing P193L, R252C or F382V receptor and those expressing WT receptor, but was significantly reduced in cells expressing the F248C mutant. No significant stimulation of glucose uptake was seen in cells expressing S323L, D707A or P1178L receptors (Figure 5).

Antibodies 83-7 and 83-14 alone stimulated glucose uptake via P193L, S323L, F382V and D707A receptors, while antibodies 18–44 and 18–146 were again less effective across the full range of mutants (Figure 5). While the magnitude of antibody-stimulated uptake was less than seen with insulin *via* WT, P193L, F248C, R252C, and F382V receptors, antibodies 83-7, 83-14 and 18–44 were more effective than insulin at stimulating glucose uptake via D707A. Dual treatment with antibodies plus insulin did not enhance glucose uptake compared to insulin alone acting *via* WT, P193L, F248C, R252C, and F382V receptors, or antibody alone when acting *via* S323L and D707A receptors.

## Discussion

Autosomal recessive insulin receptoropathies feature failure to thrive, extreme metabolic derangement, and childhood mortality, and response to therapy is poor. Longitudinal studies of genetic receptoropathies suggest a steep relationship between residual INSR function and clinical outcome: loss of 50% human INSR function, as in the parents of infants with Donohue syndrome, does not produce insulin resistance in lean people. Heterozygous dominant negative mutations produce severe insulin resistance, diagnosed peripubertally in females and later in males, and reduce receptor function to 25% or less of WT. The severe recessive receptoropathies that this study focuses on confer greater loss of function, however even with 0–25% residual function a dramatic range of phenotypes is seen, with complete or near complete loss of function producing Donohue syndrome and lethality in infancy, and less extreme loss of function producing Rabson Mendenhall syndrome, in which survival to the second or third decades is normal. Collectively these observations suggest that even modest improvements in receptor signaling in recessive disease may have decisive clinical benefit.

Many pathogenic INSR mutations are known, including more than 100 missense mutations. A subset are expressed at the cell surface but show impaired insulin binding, signal transduction, or internalization and recycling. This subset may be amenable to non conventional activation by antibody. Proof of this principle came from demonstration that two bivalent antibodies stimulated kinase activity of a single solubilized mutant receptor (F382V*(7)*), and, independently, that one bivalent antibody increased glycogen synthesis acting via a mutant receptor expressed in intact cells (S232L*(18)*). We extend these limited previous findings with systematic characterization of multiple receptor mutants and antibodies in two cellular systems, with several readouts of physiologically important responses including adipocyte glucose uptake.

One of the mutants assessed, F248C, is novel. It lies close to the well-studied R252C mutant, which is expressed but exhibits impaired internalization after insulin exposure. F248C shows only minor reduction in cell surface expression, but insulin-stimulated receptor autophosphorylation and downstream signaling are severely impaired. Across known mutants our data generally agree with prior studies. Assay of receptor autophosphorylation in CHO cells using immunocapture of myc-tagged receptor prior to immunoassay demonstrated signaling defects more clearly than phosphotyrosine immunoblotting in the 3T3-L1 overexpression system. This is likely due to the inherently greater dynamic range of immunoassay allied to use of a generic anti-phosphotyrosine antibody, with different degrees of receptor overexpression between the two models possibly also contributing.

We confirmed that S323L and F382V receptors can be partially activated by antibodies, and extended these observations to a wider range of mutants. Previous studies suggest that receptor activation by antibody depends on receptor cross-linking rather than reaction at specific epitopes*(20)*. Consistent with this, two of the antibodies we employed, 83-7 and 83-14 are both effective despite recognizing different epitopes and having different effects on insulin binding. Antibodies 18–44 and 18–146 consistently elicited much smaller responses, although 18–44 has previously been found to exert insulin-like activity on primary human adipocytes*(21)*. Differences among antibodies are likely to reflect differences of affinity and/or steric constraints on cross-linking receptors in intact cells.

The mutants showing the largest antibody response were S323L and D707A, both being activated by antibodies similarly to WT receptor, and to a greater extent than by insulin. Such mutants with ‘pure’ insulin binding defects are particularly attractive therapeutic targets. Other mutants studied in both cell systems (P193L, F248C, R252C, and F382V) all showed some activation of AKT, GSK3, AS160 and glucose uptake by antibodies. In these cases responses were less than for WT receptor or those induced by insulin although it is unclear that the conditions studied optimally model the *in vivo* environment, and receptor overexpression is likely to have mitigated some functional defects. Testing the therapeutic potential of antibodies against such mutants is warranted in vivo, where antibody signalling may be prolonged compared to insulin signalling due to slower receptor internalisation. Indeed, a previously studied anti insulin receptor antibody showed markedly greater hypoglycemic effects *in vivo* in wild type animals than had been apparent in cell culture models *(14)*.

Antibodies would be a particularly appealing therapeutic proposition were they to exhibit synergy with insulin in receptor stimulation, amplifying insulin action rather than simply imposing a tonic signal. The current studies have not addressed this in detail, although suggestive evidence for synergic stimulation of WT receptor and some mutant receptors is seen. This was not mirrored by detectable synergistic activation of downstream signaling or metabolic endpoints, possibly because maximal downstream signaling requires only submaximal receptor autophosphorylation. It remains possible that insulin-antibody synergy does exist but was obscured under the conditions of the experiments undertaken, which pragmatically employed relatively high concentrations of insulin and antibody.

Early cellular studies of antibody-induced insulin receptor activation were interpreted as suggesting that antibodies elicit greater downstream responses than expected from low levels of receptor autophosphorylation*(17, 34–36)*. These observations were later argued to have a methodological basis, hinging on lower sensitivity in detecting tyrosine phosphorylation than downstream signaling *(37, 38)*. This is in part because signal amplification is an inherent property of signal transduction cascades. Our observation of apparent “escape” from signaling inhibition in the face of efficient *Insr* knockdown in 3T3-L1 adipocytes supports this contention, as activation of residual receptors is undetectable directly but is observable downstream due to signal amplification.

Importantly, receptor activation by antibodies leads to selective AKT phosphorylation, which is critical for metabolic actions of insulin, with little or no ERK phosphorylation. As activation of the RAS/RAF/MEK/ERK pathway is mitogenic, this is an encouraging property of antibodies for translational purposes, suggesting that they may exert metabolic benefits without undue mitogenic activity. Similar dissociation between activation of AKT and ERK has also been observed following INSR activation by the peptide ligand S597*(39)* and in previous studies with anti-receptor antibodies *(40)*. The mechanism underlying such pathway-specific activation is poorly understood, although IRS proteins may be preferentially phosphorylated by plasma membrane-associated receptor*(41, 42)*, whereas receptor internalization is required for full ERK activation*(42, 43)*. This suggests that differential signaling may relate to cell surface retention of antibody-bound receptors.

In summary, multiple monoclonal antibodies can bind and activate a range of mutated cell surface insulin receptors to a potentially highly clinically significant degree. Prior experience in wild type animals*(14)* and theoretical considerations argue that effects of anti-INSR antibodies in vivo may be greater than those in cells, and so further studies of mutant human INSR in animal models are warranted. The murine antibodies studied are particularly amenable to this, as their effect will not be attenuated by host immune responses to human antibodies. Later translation to humans will require humanisation of the antibodies, however many current therapeutic antibodies are humanised or chimeric versions of murine antibodies*(9)*, and moreover humanisation has already been reported for a different purpose for 83-14, with no deleterious consequences for wild type receptor binding*(44, 45)*. This suggests a practical route to future testing of anti-INSR antibodies in severely unwell infants with recessive lethal receptoropathy where the potential clinical benefits are transformational, and also raises the prospect of later evaluation of antibodies as novel ultralong-acting insulin mimetics in more common forms of diabetes, especially where control is labile, or in settings where regular injection of insulin is challenging.

## Acknowledgments

Funding was from an Open Funding grant from the Diabetes Research and Wellness Foundation (to GVB), and a project grant from Diabetes UK (to RKS). RKS is funded by the Wellcome Trust (WT098498), and core support was provided by the Medical Research Council [MRC_MC_UU_12012/5] and the United Kingdom National Institute for Health Research (NIHR) Cambridge Biomedical Research Centre. We thank Cornelia Gewert and Danielle Newby for technical support.

## Author Contributions

GVB, KS and RKS designed the experiments. GVB performed the experiments. GVB, KS, and RKS analysed the data. GVB, KS, and RKS and wrote the manuscript. RKS is the guarantor of this work and, as such, had full access to all the data in the study and takes responsibility for the integrity of the data and the accuracy of the data analysis.

## Prior Presentation

This study was presented as a poster at the Diabetes UK Professional Conference, Manchester, UK in March 2017 and at the International Symposium on Insulin Receptor and Insulin Action, Nice, France in April 2017.

